# Distinct neural components of visually guided grasping during planning and execution

**DOI:** 10.1101/2023.01.22.525053

**Authors:** Lina K. Klein, Guido Maiello, Kevin M. Stubbs, Daria Proklova, Juan Chen, Vivian C. Paulun, Jody C. Culham, Roland W. Fleming

**Affiliations:** Department of Experimental Psychology, Justus Liebig University Giessen, Germany; Department of Psychology, University of Western Ontario, London, ON, Canada; Center for the Study of Applied Psychology, Guangdong Key Laboratory of Mental Health and Cognitive Science, and the School of Psychology, South China Normal University, Guangzhou, Guangdong Province, 510631, China; Key Laboratory of Brain, Cognition and Education Sciences (South China Normal University), Ministry of Education; McGovern Institute for Brain Research, Massachusetts Institute of Technology, Cambridge, MA, USA; Department of Brain and Cognitive Sciences, Massachusetts Institute of Technology, Cambridge, MA, USA; Center for Mind, Brain and Behavior, University of Marburg and Justus Liebig University Giessen, Germany

**Keywords:** Grasping, functional Magnetic Resonance Imaging, Visual Grasp Selection, Grasp Axis, Grasp Size, Object Mass, Material Perception

## Abstract

Selecting suitable grasps on three-dimensional objects is a challenging visuomotor computation, which involves combining information about an object (e.g., its shape, size, and mass) with information about the actor’s body (e.g., the optimal grasp aperture and hand posture for comfortable manipulation). Here we used functional magnetic resonance imaging to investigate brain networks associated with these distinct aspects during grasp planning and execution. Human participants viewed and then executed preselected grasps on L-shaped objects made of wood and/or brass. By leveraging a computational approach that accurately predicts human grasp locations, we selected grasp points that disentangled the role of multiple grasp-relevant factors: grasp axis, grasp size, and object mass. Representational Similarity Analysis revealed that grasp axis was encoded along dorsal-stream regions during grasp planning. Grasp size was first encoded in ventral-stream areas during grasp planning, then in premotor regions during grasp execution. Object mass was encoded in ventral-stream and (pre)motor regions only during grasp execution. Premotor regions further encoded visual predictions of grasp comfort, whereas the ventral stream encoded grasp comfort during execution, suggesting its involvement in haptic evaluation. These shifts in neural representations thus capture the sensorimotor transformations that allow humans to grasp objects.

**Significance Statement:** Grasping requires integrating object properties with constraints on hand and arm postures. Using a computational approach that accurately predicts human grasp locations by combining such constraints, we selected grasps on objects that disentangled the relative contributions of object mass, grasp size, and grasp axis during grasp planning and execution in a neuroimaging study. Our findings reveal a greater role of dorsal-stream visuomotor areas during grasp planning, and surprisingly, increasing ventral stream engagement during execution. We propose that during planning, visuomotor representations initially encode grasp axis and size. Perceptual representations of object material properties become more relevant instead as the hand approaches the object and motor programs are refined with estimates of the grip forces required to successfully lift the object.

## Introduction

Grasping is one of the most frequent and essential everyday actions performed by humans and other primates (Betti et al., 2021), yet planning effective grasps is computationally challenging. Successful grasping requires identifying object properties including shape, orientation and mass, and considering how these interact with the capabilities of our hands (Fabbri et al., 2016; Maiello et al., 2019, 2021; Klein, Maiello et al., 2020). Whether an object is large or small, heavy or light, determines how wide we open our hands to grasp it and how much force we apply to lift it (Johansson and Westling, 1988; Cesari and Newell, 1999). Such grasp-relevant object properties, including weight, mass distribution, and surface friction can often be inferred visually before initiating actions (Fleming, 2017; Klein et al., 2021).

A recent computational model accurately predicts precision-grip grasp locations on 3D objects of varying shape and non-uniform mass (Klein, Maiello et al., 2020). The model combines multiple constraints related to properties of the object and the effector, such as the torque associated with different grasps and the actor’s natural grasp axis. However, it remains unclear which brain networks are involved in computing specific grasping constraints. Moreover, it is unknown whether all constraints are estimated during grasp planning (i.e., before action initiation; Gallivan et al., 2013, 2019) or whether some aspects are computed during action execution, allowing the actor to refine grasp parameters on-line before or during contact with the object. Here, we ask how information gets combined to evaluate and then execute grasps. While many previous studies have investigated the effects of individual attributes, during either grasp planning or execution, here we consider how multiple factors combine, and compare both planning and execution.

Previous studies show that grasp-relevant representations are distributed across ventral and dorsal visual processing streams. Shape is represented throughout both streams (Sereno et al., 2002; Orban et al., 2006; Konen and Kastner, 2008; Orban, 2011), with dorsal representations emphasizing information required for grasp planning (Srivastava et al., 2009). For example, dorsomedial area V6A—located in human superior parieto-occipital cortex (SPOC)—is involved in selecting hand orientation given object shape (Fattori et al., 2004, 2009, 2010; Monaco et al., 2011). Visual representations of material properties—also crucial for grasping—have been identified predominantly in ventral regions such as lateral occipital cortex (LOC), the posterior fusiform sulcus (pFS), and parahippocampal place area (PPA; Cant and Goodale, 2011; Hiramatsu et al., 2011; Gallivan et al., 2014; Goda et al., 2014, 2016). Brain regions that transform these disparate visual representations into appropriate motor codes include Anterior Intraparietal Sulcus (aIPS), Ventral Premotor Cortex (PMv), Dorsal Premotor Cortex (PMd), and primary motor cortex (M1). Primate neurophysiology suggests that PMv (primate Area F5) encodes grip configuration (Murata et al., 1997; Raos et al., 2006; Theys et al., 2012), while PMd (primate Area F2) encodes grip/wrist orientation (Raos et al., 2004). Both regions exhibit strong connections with aIPS, which could play a key role in linking visual representations— including those in ventral stream regions (Borra et al., 2008)—to motor commands sent to the hand through M1 (Murata et al., 2000; Janssen and Scherberger, 2015).

How information flows and is combined across this complex network of brain regions is far from understood. We therefore sought to identify cortical regions associated with distinct components of grasping and tested their relative importance during grasp planning and execution. To disentangle grasping constraints, we used our model (Maiello et al., 2021) to select grasps that placed different constraints in conflict. For example, a selected grasp could be near optimal in terms of the required hand axis, but sub-optimal in terms of grasp aperture. We then measured functional magnetic resonance imaging (fMRI) blood-oxygen-level-dependent (BOLD) activity, during planning and execution of these preselected grasps. Combining this model-guided approach with representational similarity analysis (RSA; Kriegeskorte, 2008) let us tease apart the relative contributions of object mass, grasp size, and grasp axis, at different stages of grasping.

## Results

Participants in a 3-Tesla MRI scanner were presented with physical 3D objects on which predefined grasp locations were shown (**Figure 1A**). On each trial, participants first planned how to grasp the objects (planning phase, **Figure 1B**) and then executed the grasps (execution phase). We designed objects and grasp locations to produce a set of nine distinct conditions (**Figure 1C**) that would differentiate three components of grasping: the grasp axis (i.e., orientation), the grasp size (i.e., the grip aperture), and object mass. By computing pairwise distances between all conditions for each of these grasp-relevant dimensions, we constructed one representational dissimilarity matrix (RDM) for each component (**Figure 1D-F**)—these were uncorrelated across conditions. In each brain region of interest (ROI) tested in the study (**Figure 1H**), brain-activity patterns elicited by each condition were compared to each other via Pearson correlation to construct brain RDMs. **Figure 1G** shows one such RDM computed from brain region PMv for one example participant during the planning phase. In this participant, this area appeared to strongly encode grasp axis.

**Figure 1.**
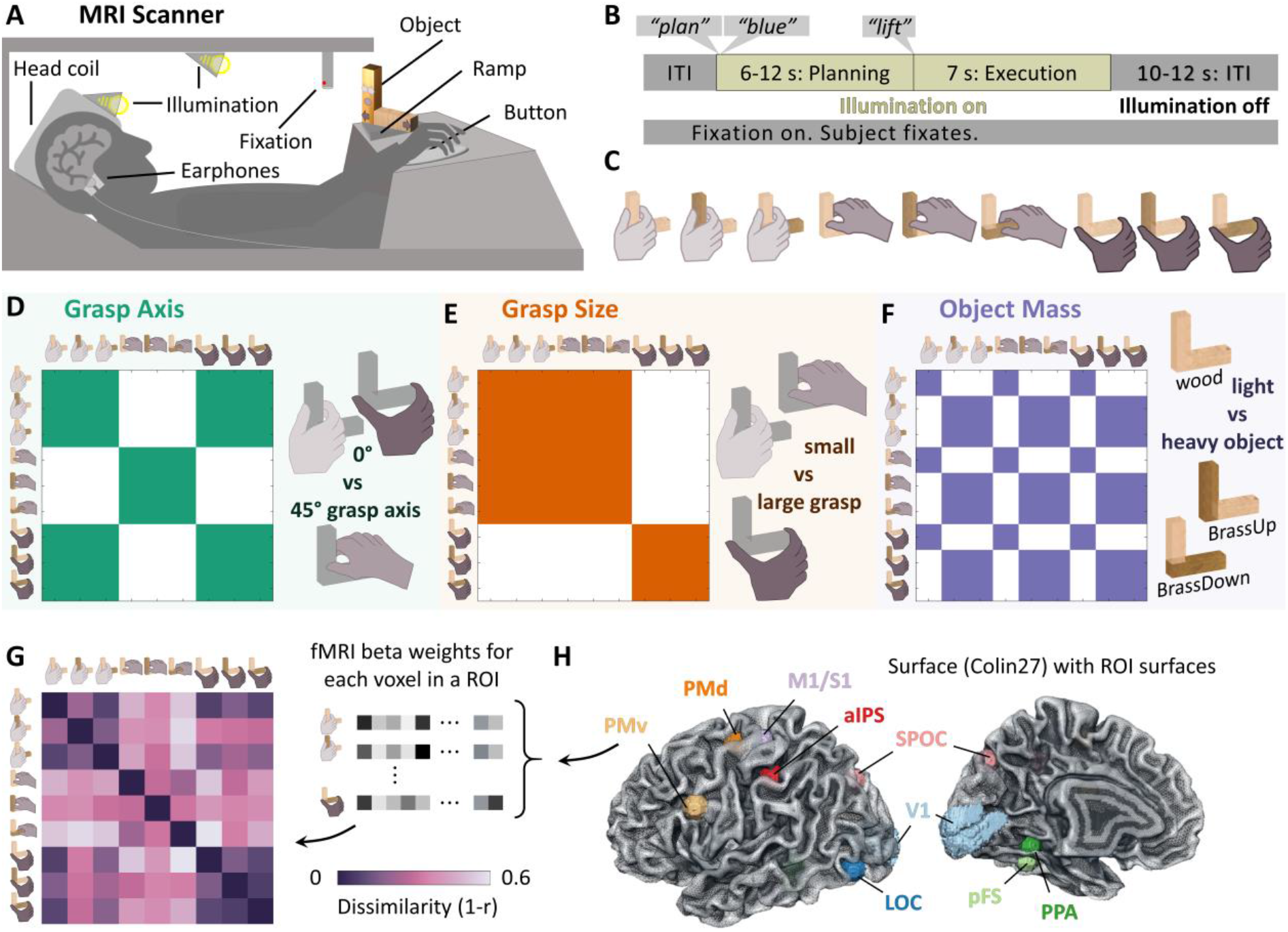
Study design. **(A)** Participants in the MRI scanner were cued to grasp 3D objects at specific locations. **(B)** Sequence of events for one example trial during which participants were instructed to grasp the object at the predefined location marked by different colour dots or arrows. **(C)** Preselected grasps on stimulus objects of wood and brass produced nine distinct conditions designed to differentiate three components of grasping using RSA. **(D-F)** RDMs for grasp axis, grasp size, and object mass. Coloured cells represent condition pairs with zero dissimilarity, white cells represent maximum dissimilarity. **(G)** An example RDM computed from fMRI BOLD activity patterns in region PMv of one participant during the planning phase. Note the strong similarity to the grasp axis RDM in panel D. (**H**) Visualization of the selected ROIs within the Colin27 template brain. All ROIs except V1 were built as spheres centred on coordinates recovered from neurosynth.org. V1 coordinates were taken from the (Wang et al., 2015) atlas. Note that surface-rendering is for presentation purposes only as data were analysed in volumetric space and no cortex-based alignment was performed.

### How grasp-relevant neural representations develop across the grasp network

**Figure 2A** shows average neural RDMs computed throughout the network of visuomotor brain regions we investigated. ROIs were selected from the literature as regions most likely specialized in the components of visually guided grasping investigated in our study. We included primary visual cortex, V1, as the first stage of cortical visual processing. Areas LOC, pFS, and PPA within the ventral visual stream (occipitotemporal cortex) were included as they are known to process visual shape and material appearance (Cant and Goodale, 2011; Hiramatsu et al., 2011; Gallivan et al., 2014; Goda et al., 2014, 2016), and could thus be involved in estimating object mass. Areas SPOC, aIPS, PMv, and PMd within the dorsal visual stream (occipitoparietal and premotor cortex) were included as they are thought to transform visual estimates of shape and orientation into motor representations (Janssen and Scherberger, 2015). Primary motor and somatosensory area (M1/S1, in the central sulcus) was included as the final stage of cortical sensorimotor processing. The patterns of correlations between model and neural RDMs across participants and ROIs (**Figure 2B-G**) reveal which information was encoded across these visuomotor regions during grasp planning and execution phases.

**Figure 2.**
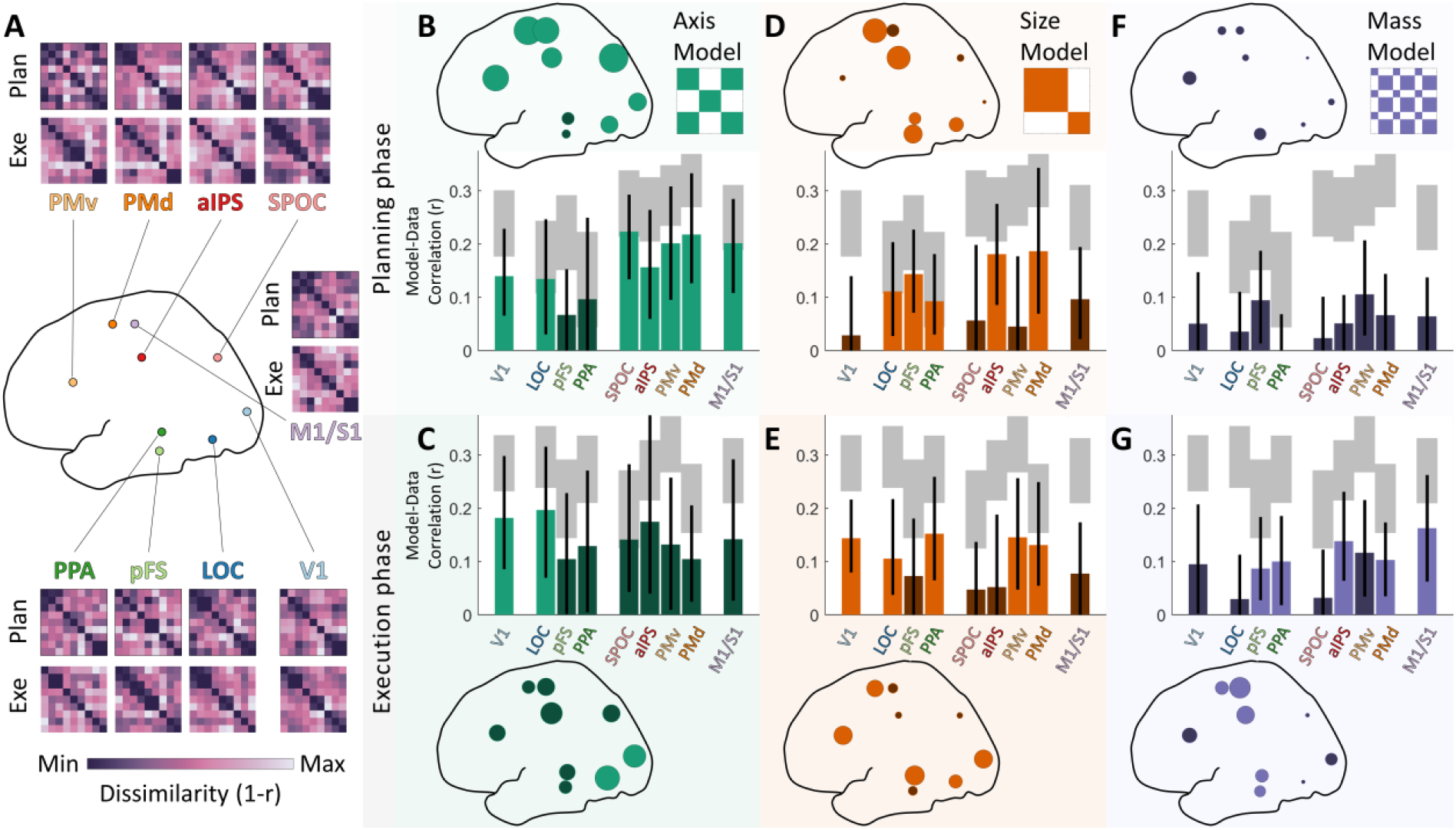
RSA results. **(A)** Mean neural RDMs computed in the nine ROIs included in the study. For visualization purposes only, RDMs within each region are first averaged across participants and then normalized to the full range of the LUT. **(B-G)** Correlations between model and neural RDMs in each brain ROI during planning (top, B,D,F) and execution phases (bottom, C,E,G). In bar graphs, grey shaded regions represent the noise ceiling for each ROI. Bars are means, error bars represent 95% bootstrapped confidence intervals. The same data are represented topographically as dots scaled proportionally to the mean correlation in each region. Bright colours represent significant positive correlations (p<.05 with FDR correction); correlations shown in dark colours are not statistically significant.

### Grasp axis encoding in visuomotor regions during grasp planning

**Figure 2B,C** shows that neural representations in V1 and ventral region LOC were significantly correlated with grasp axis during both grasp planning and execution phases. In contrast, representations in ventral areas pFS and PPA were never significantly correlated with grasp axis. Further, grasp axis was significantly correlated with neural representations across all dorsal areas (SPOC, aIPS, PMv, PMD), as well as M1/S1, but only during grasp planning. Dorsal and motor areas thus robustly encoded the orientation of the hand when preparing to grasp objects, suggesting that the hand-wrist axis was among the first components of the action computed across these regions.

### Grasp size was encoded across both visual streams during grasp planning and execution

During the planning phase (**Figure 2D**), grasp size significantly correlated with neural representations in all ventral areas (LOC, pFS, PPA), and with representations in dorsal regions aIPS and PMd. During the execution phase (**Figure 2E**), grasp size remained significantly correlated with neural representations in ventral areas LOC and PPA, but not pFS. In the dorsal stream during the execution phase, grasp size remained significantly correlated with neural representations in PMd but not aIPS, and became significantly correlated with representations in PMv. Neural representations in early visual area V1 were significantly correlated with grasp size only in the execution phase, but not during planning. Finally, neural representations in sensorimotor area M1/S1 were never significantly correlated with grasp size. Thus, different ventral and dorsal areas encoded grasp size at different time points. These data suggest that ventral regions may have been initially involved in computing grasp size and might have relayed this information (e.g., through aIPS) to the premotor regions tasked with generating the motor codes to adjust the distance between fingertips during the execution phase.

### Object mass was encoded across dorsal and ventral streams and in motor areas, but only during grasp execution

During the planning phase (**Figure 2F**), none of the investigated ROIs exhibited any activity that was significantly correlated with object mass. Conversely, during the execution phase (**Figure 2G**), object mass significantly correlated with representations in ventral areas pFS and PPA, dorsal areas aIPS and PMd, and sensorimotor area M1/S1. Object mass was thus encoded in the later stages of grasping. One possible interpretation is that this occurred when the hand was approaching the object and was preparing to apply appropriate forces at the fingertips. Alternatively, it could be due to sensory feedback about slippage once the object was lifted.

### Representational similarities within the grasp network

We took the RDMs generated for each of the nine ROIs (**Figure 2**) and correlated them with one another to reveal inter-ROI similarity relationships. **Figure 3** summarizes the resulting second-order similarity relationships, both within and between planning and execution phases.

**Figure 3.**
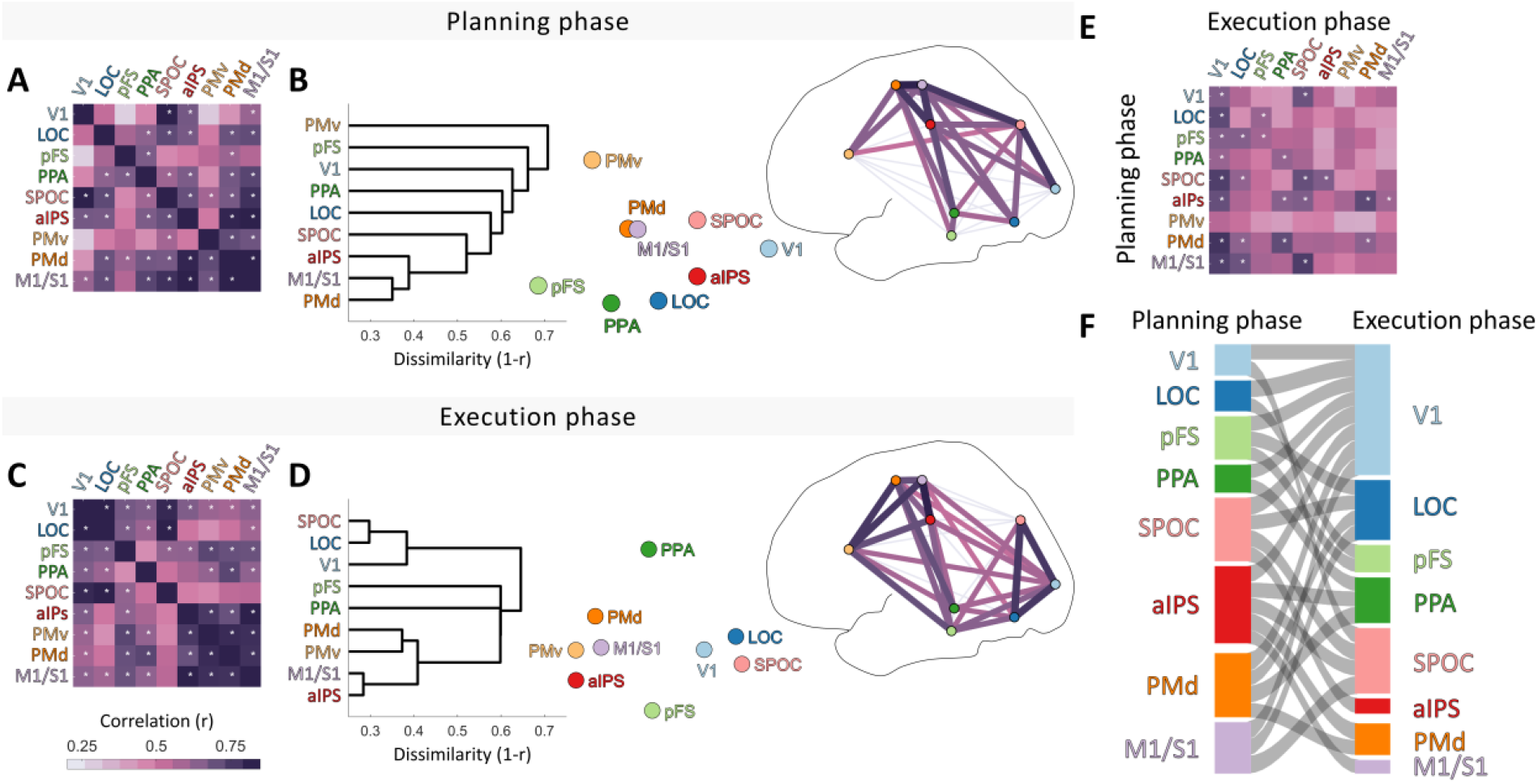
The representational structure of grasping. **(A)** Matrix showing correlations of data RDMs between regions during the planning phase. White asterisks represent significant correlations (p<.05 with Bonferroni correction). **(B)** The same data in A are shown through hierarchical clustering and 2D multidimensional scaling, and significant correlations are shown topographically. **(C**,**D)** As in A, except for the planning phase. **(E)** Correlations between ROIs across planning and execution phases. **(F)** Sankey diagram depicting significant correlations from E.

We find that neural representations were significantly correlated across many selected ROIs during both grasp planning (**Figure 3A**) and execution (**Figure 3C**). Of particular note is that during the planning phase, dorsal regions tended to correlate more strongly with one another, while during the execution phase, ventral regions showed more correlated representations. This is revealed by visualising the inter-ROI similarities arranged topographically within a schematic brain (**Figure 3B and 3D**), with the darkness of connecting lines between ROIs proportional to the correlations between their corresponding RDMs.

During planning (**Figure 3B**), the strongest correlations were between M1/S1, PMd and aIPS; between V1 and SPOC; and to a lesser extent between SPOC and M1/S1. In the execution phase (**Figure 3D**) the similarities among brain regions formed two main clusters. One cluster of visual regions was formed by V1, SPOC, and LOC. The second cluster comprised aIPS, premotor areas PMv and PMd, and M1/S1. MDS and topographical plots highlight how these two clusters appeared to share representational content predominantly through ventral stream regions pFS and PPA.

### Shared representations across planning and execution phases

Neural representation patterns were also partly correlated across grasp planning and execution phases (**Figure 3E,F**). Notably, aIPS representations during the planning phase were significantly correlated with representational patterns in ventral (PPA), dorsal (SPOC, PMd), and sensorimotor (M1/S1) regions during the execution phase. This suggests that aIPS may play a key role in linking grasp planning to execution. Further, neural representation patterns in nearly all ROIs (except PMv) during the planning phase were correlated with representations in V1 during the execution phase, and representations in PFs, SPOC, PMd, and M1/S1 during action planning were correlated with LOC representations during action execution. We speculate that this might reflect mental simulation, prediction, and feedback mechanisms at play (see **Discussion**).

### Grasp comfort

We recently demonstrated that humans can visually assess which grasp is best among competing options and can refine these judgements by executing competing grasps (Maiello et al., 2021). These visual predictions and haptic evaluations of grasp comfort were well captured by our multi-factorial model (Klein, Maiello et al., 2020), suggesting they may play a role in grasp selection. We thus wondered whether we could identify, within the grasp network investigated here, brain regions that encoded visual predictions and haptic evaluations of grasp comfort. To this end, once an imaging session was completed, we asked participants (while still lying in the scanner) to execute once more each of the nine grasps and rate how comfortable each felt on a scale of 1 to 10. Comfort ratings were consistent across participants (**Figure 4A**). Comfort was slightly modulated by grasp axis (**Figure 4B**, t(20)=3.3, p=.0037) and was not modulated by grasp size (**Figure 4C**, t(20)=0.89, p=.39). The factor that most affected grasp comfort was object mass, with heavy objects being consistently rated as less comfortable than light objects (**Figure 4D**, t(20)=8.1, p<.001). This was also evident when we computed RDMs from comfort ratings (**Figure 4E**) and found that these were significantly correlated with the model RDM for object mass (p<.001) but not with RDMs for grasp axis (p=.54) or grasp size (p=.83) (**Figure 4F**).

**Figure 4.**
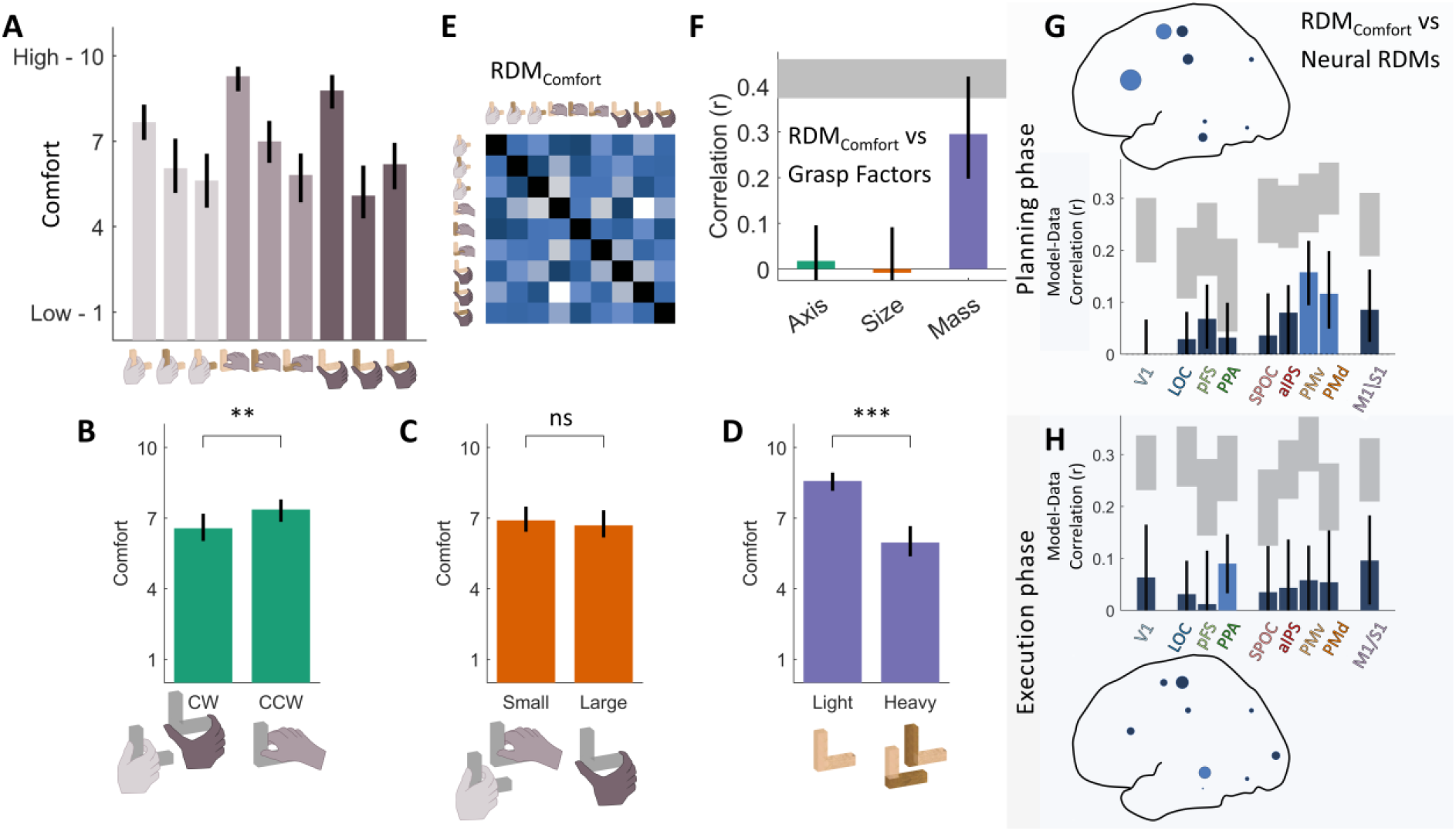
Grasp comfort. **(A)** Average grasp comfort ratings for each grasp condition in the fMRI experiment. **(B**,**C**,**D)** Grasp comfort ratings averaged across (B) grasp axis, (C) grasp size, and (D) object mass. **(E)** Average RDM computed from participant comfort ratings. **(F)** Correlations between grasp comfort and model RDMs. **(G**,**F)** Correlations between grasp comfort and neural RDMs in each brain ROI during planning (top, G) and execution phases (bottom, H). In bar graphs, grey shaded regions represent the noise ceiling for each ROI. Bright blue bars represent significant positive correlations (p<.05 with FDR correction); correlations shown in dark blue are not statistically significant. The same data are represented topographically as dots scaled proportionally to the mean correlation in each region. Across figure panels, bars are means, error bars represent 95% bootstrapped confidence intervals. **p<0.01, ***p<0.001

### Neural representations of grasp comfort were present during both grasp planning and execution phases

To identify brain regions that encoded grasp comfort, we next correlated neural RDMs with the average RDM derived from participant comfort ratings. Neural representations in premotor regions PMv and PMd were significantly correlated with grasp comfort during grasp planning (**Figure 4G**). During the execution phase instead, grasp comfort correlated with neural representations in ventral stream region PPA (**Figure 4H**). This suggests that dorsal premotor regions encoded the visually predicted comfort of planned grasps (which in our conditions was primarily related to the object mass). Area PPA instead encoded comfort during the execution phase, and might thus be involved in the haptic evaluation of grasp comfort, or some other representation of material properties that correlate with comfort.

## Discussion

Our results show that different brain regions within the two visual streams represent distinct determinants of grasping, including grip axis, grip size, and object mass; moreover, the coding of these attributes differed between grasp planning and execution. Most regions represented multiple factors at different stages. For example, aIPS activity correlated with both grasp axis and grasp size during planning, and with object mass during execution. We found that grasp axis, which is adjusted at the very beginning of reach-to-grasp movements (Cuijpers et al., 2004), was predominantly encoded across dorsal regions during grasp planning. Grasp size, which is adjusted throughout reach-to-grasp movements (Cuijpers et al., 2004), was encoded in different sets of ventral and dorsal regions during grasp planning and execution. Object mass, which gains relevance when applying forces at the fingertips upon hand-object contact (Johansson and Westling, 1988; Johansson and Flanagan, 2009), was instead encoded across ventral, dorsal and motor regions during grasp execution.

### Shift from dorsal-to ventral-stream regions between planning and execution

In the broadest of terms, our analysis revealed an overall shift—in terms of representational similarity—from dorsal sensory and motor regions during the planning phase (**Figure 3AB**) to more ventral regions during execution (Figure 3CD). Specifically, during planning the most similar representations were between V1 and SPOC, SPOC and M1/S1, and between M1/S1, PMd and aIPS, tracing an arc along the dorsal stream to frontal motor areas. SPOC is associated with representations of grasp axis (Monaco et al, 2011), as is parieto-occipital area V6A in the macaque, which together with V6 is thought to be the macaque homologue of human SPOC (Fattori et al., 2004, 2009, 2010; Pitzalis et al., 2013). The SPOC complex serves as a key node in the dorsal visual stream involved in the early stages of reach to grasp movements (Rizzolatti and Matelli, 2003). It is thus interesting to speculate that our findings likely represent the progressive transformation of grasp-relevant sensory representations of the object into explicit motor plans along the dorsal processing hierarchy. In contrast, along the ventral stream, individual ROIs (V1, LOC, PPA, pFS) shared similar representations with dorsal sensorimotor areas (particularly aIPS, M1/S1 and PMd), but only weak or no correlation with one another (or with PMv). During planning there was no visual movement to drive common responses and it seems reasonable to assume that different ROIs extracted distinct aspects of the stimulus, leading to the rather weak correlations.

During action execution, the picture changed dramatically. Representations in the dorsal stream became more independent from one another. Notably, the high similarity between SPOC representations and the more frontal motor regions (M1/S1, aIPS, PMd and PMv) almost disappeared, to be replaced with a stronger correlation with ventral shape-perception area LOC. At the same time, representational correlations between ventral visual regions V1, LOC, PPA and pFS, as well as their correlations with PMv increased. This may partly be due to the salient visual consequences of the participant’s own actions providing a common source of variance across the regions. It is interesting to speculate that the overall shift from similar dorsal to similar ventral representations reflects a shift from the extraction of action relevant visual information during planning to monitoring object properties to assess the need for corrections during the action execution phase.

One of the more striking findings from representational similarity analysis (**Figure 3E,F**) is that activity in V1 during execution correlated with representations in a slew of high visual and sensorimotor areas during the planning phase (this is visible as the column of dark values below V1 in **Figure 3E**, and as the large and dense pattern of connections towards V1 in the Sankey plot in **Figure 3F**).

We speculate that the shift in representations between planning and execution might reflect a role of mental simulation in grasp planning and subsequent comparison to the sensory evidence during execution. During the planning phase, participants may be utilizing visual information to compute and compare forward models of potential grip choices (Wolpert and Flanagan, 2001; Cisek and Kalaska, 2010), and possibly mentally simulating potential grasps (Jeannerod, 1995; Jeannerod and Decety, 1995). These simulations could be used to generate motor plans and sensory predictions. Sensory predictions could then be compared to visual, tactile, and proprioceptive inputs during the grasping phase, to facilitate online movement corrections and evaluate the success of the generated motor plan (Desmurget and Grafton, 2000; Wolpert and Ghahramani, 2000; Wolpert et al., 2011). This possibility is supported by recent work showing that planned actions can be decoded from activity in V1 and LOC before movement onset (Gallivan et al., 2013, 2019; Gutteling et al., 2015; Monaco et al., 2020), and that V1 and LOC are re-recruited when performing delayed actions toward remembered objects (Singhal et al., 2013).

### Effects of grasp comfort

Grasp comfort was moderately correlated with object mass (*r* ∼ 0.3) but not grip axis nor grip size, suggesting that other factors were also affected comfort (perhaps even more so than usual because of the movement constraints in the scanner). Grasp comfort was significantly correlated PPA activation during execution, perhaps related to a role for PPA in also coding object mass during execution. More interestingly, activation patterns in premotor cortex (PMv and PMd) were correlated with grasp comfort during planning even though no regions significantly represented object mass during planning. These results corroborate earlier results implicating premotor cortex in grip selection based on orientation (Martin et al., 2011; Wood et al., 2017) and extend the findings to a broader range of factors and to multivariate representations.

### Limitations and future directions

One notable finding of our study is that object mass is encoded in sensorimotor regions during action execution. This is understandable, as information about object mass is required to modulate grip and lift forces. However, we have previously demonstrated that mass and mass distribution also play an important role in selecting where to grasp an object (Klein, Maiello et al., 2020). It is thus reasonable to expect processing of object material and mass also during planning, which we did not observe. However, in our study, grasps were preselected. As a result, participants did not need to process an object’s material properties to select appropriate grasp locations. In order to investigate the role of visual material representations in grasp selection, future research could use our computational framework (Klein, Maiello et al., 2020; Maiello et al., 2021) to identify objects that produce distinct grasp patterns, rather than constraining participants to predefined grasp locations. Conditions that require visual processing of object material properties to select appropriate grasp locations would then reveal whether the same or different sensorimotor regions process object mass during grasp planning and execution. However, such designs would require disentangling activity related to representing shape *per se* from activity related to grasp selection and execution.

One factor which is known to be important for grasp selection and execution is grip torque, i.e., the tendency of an object to rotate under gravity when grasped away from its centre of mass (Goodale et al., 1994; Lederman and Wing, 2003; Eastough and Edwards, 2006; Lukos et al., 2007; Paulun et al., 2016). While torque is directly related to object mass, it is possible to select different grasps on the same object which produce substantially different torques (Maiello et al., 2021). Since grasps with high torque require greater forces at the fingertips to maintain an object level, humans tend to avoid such high-torque grasps (Klein, Maiello et al., 2020). We originally designed our stimuli in the hope of dissociating torque from object mass. Unfortunately, in pilot testing we observed that certain object and grip configurations in the magnetic field of the MRI scanner produced eddy currents in the brass portions of our stimuli. These currents caused unexpected magnetic forces to act on the stimuli, which in turn altered fingertip forces required to grasp and manipulate the objects. To avoid the occurrence of such eddy currents in our experiment, we decided to forgo conditions differentiating the effects of object mass from those of grip torques. By employing nonconductive materials, in future work our approach could be extended to test whether grasp-relevant torque computations occur in the same visuomotor regions responsible for estimating object material and shape. While previous studies have investigated material and shape largely independently, one intriguing question for future research is how material and shape are combined to assess the distribution of materials and the consequences of mass distribution on torque and grip selection.

## Conclusions

Taken together, our results extend previous behavioral and modelling findings about how participants select optimal grasp based on myriad constraints (Klein, Maiello et al., 2020) to reveal the neural underpinnings of this process. Results show that distinct factors – grip orientation, grip size, and object mass – are each represented differently. Moreover, these representations change between grasp planning and execution. Representations during planning rely relatively more heavily on the dorsal ventral stream, while those during execution rely relatively more heavily on the ventral visual stream. Though surprising, this transition can be explained by a transition from grip selection during planning to monitoring of sensory feedback during grasping execution.

## Materials and Methods

### Participants

Analyses utilized data from 21 participants (13 female, mean [range] age: 25.5 [18-33]) recruited from the University of Western Ontario. Data from two additional participants were excluded due to excessive head motion. All participants had normal or corrected-to-normal vision and were fully right-handed as measured by the Edinburgh Handedness Inventory. Informed consent was given prior to the experiment. The study was approved by the Health Sciences Research Ethics Board at the University of Western Ontario and followed the principles in the sixth revision of the Declaration of Helsinki (2008). Participants were instructed on how to perform the experimental task before entering the MRI room, yet remained naïve with respect to the study’s hypotheses. All participants were financially compensated at a rate of C$25/hour.

### Setup

A schematic of our setup is shown in **Figure 1A**. Each participant lay supine inside the MRI scanner with their head placed in a head coil tilted by ∼30° to allow direct viewing of real stimulus objects placed in front of them. Below the head we positioned the bottom 20 channels of a 32-channel head coil and we suspended a 4-channel flex coil via loc-line (Lockwood Products, Inc.) over the forehead. A black wooden platform, placed above a participant’s hip, enabled the presentation of real objects that participants were required to grasp, lift, and set back down using their right hand. The platform’s flat surface was tilted by ∼15° towards a participant in order to maximize comfort and visibility. Objects were placed on a black cardboard target ramp (**Figure 1A**: “Ramp”, dimensions: 15 × 5 × 13 cm) on top of the platform that created a level surface which prevented objects from tipping over. The objects’ exact placement was adjusted such that all required movements were possible and comfortable. Between trials, a participant’s right hand rested on a button at a start position on the table’s lower right side. The button monitored movement start and end times. A participant’s upper right arm was strapped to their upper body and the MRI table using a hemi-cylindrical brace (not displayed in Figure 1A). This prevented shoulder and head movements, thus minimizing movement artefacts while enabling reach-to-grasp movements through elbow and wrist rotations. A small red LED fixation target was placed above and at a slightly closer depth location than the object to control for eye movements. Participants were required to maintain fixation on this target at all times during scanning. An MR-compatible camera was positioned on the left side of the head coil to record the participant’s actions. Videos of the runs were screened offline and trials containing errors were excluded from further analyses. A total of 22 error trials were excluded, 18 of which occurred in one run where the participant erroneously grasped the objects during the planning phase.

Two bright LEDs illuminated the workplace for the duration of the planning and execution phases of each trial, one was mounted on the head coil and the other was taped to the ceiling of the bore. Another LED was taped to the outside of the bore and was only visible to the experimenter to cue the extraction and placement of the objects. The objects were kept on a table next to the MRI-scanner, on which three LEDs cued the experimenter on which object to place inside the scanner. Participants wore MR-safe headphones through which task instructions were relayed on every trial. The LEDs and headphones were controlled by a MATLAB script on a PC that interfaced with the MRI scanner. Triggers were received from the scanner at the start of every volume acquisition. All other lights in the MRI room were turned off and any other potential light sources and windows were covered so that no other light could illuminate the participant’s workspace.

### Stimuli

Stimuli were three L-shaped objects of the same size, created from seven blocks (cubes of 2.5 cm side length). One object was constructed with seven cubes of beech wood (object weight: 67g), whereas the other two were both constructed of four brass and three wooden cubes (object weight: 557g). The two identical wood-brass objects were positioned in two different orientations, one with the brass “arm” pointing up (see **Figure 1F**: “BrassUp”), the other with the brass arm lying down (“BrassDown”). In a slow event-related fMRI design, on each trial participants directly viewed, grasped, and lifted an object placed on a platform.

### Task

Participants performed three distinct grasps per object, each grasp marked on the objects with coloured stickers during the experiment. The colours were clearly distinguishable inside the scanner and served to cue participants about which grasp to perform. Participants were instructed to perform three-digit grasps with their right hand, by placing the thumb in opposition to index and middle fingers. This grasp was similar to the precision grip grasps employed in our previous work (Maiello et al., 2019, 2021; Klein, Maiello et al., 2020; Klein et al., 2021), but ensured participants could apply sufficient grip force to lift all objects to a height of approximately 2 cm above the platform. Grasp contact locations for the index and thumb were selected in order to produce a set of uncorrelated—and thus independent—representational dissimilarity matrices (RDMs) for the three grasp factors investigated: grasp axis, grasp size, and object mass. Specifically, grasps could be rotated 45° either clockwise or counter clockwise around the vertical axis, and could require small (2.5 cm) or large (7.5 cm) grip apertures. In pilot testing we further refined the positioning of the objects and grasps within the magnetic field of the MRI scanner to avoid the forming of eddy currents within the brass parts of the objects which could hinder participants from executing the grasps. The complete set of grasp conditions is shown in **Figure 1C**.

### Procedure

#### FMRI Experiment

We employed a slow event-related fMRI design with trials spaced every 23-31 s. Participants underwent four experimental runs in which they performed each combination of 3 objects x 3 grasps twice per run in a pseudorandom order for each run (18 trials per run, 72 trials in total). The sequence of events occurring on each trial is schematized in **Figure 1B**. Prior to each trial, the experimenter was first cued on which object to place inside the scanner. The experimenter placed the object on the ramp (6-12 s before trial onset). At trial onset, the illumination LEDs turned on and over the headphones the participant heard the instruction “plan”, immediately followed by the auditory cue specifying which grasp to execute. The auditory cue was “blue”, “green”, or “red”, which corresponded to coloured stickers marking the grasp locations on the objects. The duration of the planning phase of the task was randomly selected to be 6, 8, 10, or 12 s. During this time, the participant was required to hold still and mentally prepare to grasp the object at the cued location. Once the planning phase ended, “lift” was played over headphones to cue the participant to execute the grasp. During the execution phase of the task, the participant had 7 s to reach, grasp, and lift the object straight up by approximately 2 cm, place it back down on the target ramp, and return their hand to the start position. The experimenter then removed the object and the next trial commenced. Participants were instructed about the task, familiarized themselves with the objects, and practiced the grasps outside of the MRI room for about 5 minutes prior to the experiment. Once participants were strapped into the setup, they practiced all grasps again, thus ensuring that they could comfortably grasp each object.

#### Grasp Comfort Ratings

At the end of the fMRI experiment, participants remained positioned in the scanner and performed a short rating task. Participants were asked to perform one more time each of the nine grasp conditions. For each grasp, participants verbally reported how comfortable the grasp was on a scale of 1-10 (1 being highly uncomfortable and 10 being highly comfortable). Verbal ratings were manually recorded by the experimenter.

#### Analyses

Data analyses were conducted using Brain Voyager 20.0 (BV20) and 21.4 (BV21.4) software packages (Brain Innovation, Maastricht, The Netherlands), as well as MATLAB version R2019b.

#### fMRI data acquisition

Imaging was performed using a 3-Tesla Siemens Prisma Fit MRI scanner located at the Robarts Research Institute at the University of Western Ontario (London, Ontario, Canada). Functional MRI volumes were acquired using a T2*-weighted, single-shot, gradient-echo echo-planar imaging acquisition sequence. Functional scanning parameters were: time to repetition (TR) = 1000 ms; time to echo (TE) = 30 ms; field of view = 210 × 210 mm in-plane; 48 axial 3-mm slices; voxel resolution = 3-mm isotropic; flip angle = 40°; and multi-band factor = 4. Anatomical scans were acquired using a T1-weighted MPRAGE sequence with parameters: TR = 2300 ms; field of view = 248 × 256 mm in-plane, 176 sagittal 1-mm slices; flip angle = 8°; 1-mm isotropic voxels.

#### fMRI data preprocessing

Brain imaging data were preprocessed using the BV20 Preprocessing Workflow. First, we performed Inhomogeneity Correction and extracted the brain from the skull. We then coregistered the functional images to the anatomical images, and normalized anatomical and functional data to Montreal Neurological Institute (MNI) space. Functional scans underwent motion correction and high-pass temporal filtering (to remove frequencies below 3 cycles/run). No slice scan time correction and no spatial smoothing were applied.

#### General linear model

Data were further processed with a random-effects general linear model (GLM) that included one predictor for each of the 18 conditions (3 grasp locations x 3 objects x 2 phases [planning vs. execution]) convolved with the default Brain Voyager “two-gamma” hemodynamic response function (Friston et al., 1998) and aligned to trial onset. As predictors of no interest, we included the six motion parameters (x, y, and z translations and rotations) resulting from the 3D motion correction.

#### Definition of Regions of Interest

We investigated a targeted range of regions of interest (ROIs). The locations of these ROIs are shown in **Figure 1H**; the criteria used to define the regions and their MNI coordinates are given in Table 1. ROIs were selected from the literature as regions most likely specialized in the components of visually guided grasping investigated in our study. These included primary visual cortex V1, areas LO, pFS, and PPA within the ventral visual stream (occipitotemporal cortex), areas SPOC, aIPS, PMv, PMd within the dorsal visual stream (occipitoparietal and premotor cortex), and primary sensorimotor cortex M1/S1.

**Table 1.** Regions of interest and their peak x-, y-, and z-coordinates in MNI space. Associated search term used on neurosynth.org with the number of studies the meta-analyses are based on and the extraction date (when the files were downloaded from the website). V1-coordinates were taken from (Wang et al., 2015).

| ROIs in the left hemisphere | Centre X | Centre Y | Centre Z | Search term (*neurosynth*) | Based on # of studies | Extraction date |
| --- | --- | --- | --- | --- | --- | --- |
| **V1** (primary visual) | see (Wang et al., 2015) |  |  |  |  |  |
| **LOC** (lateral occipital cortex) | -42 | -78 | -6 | *lateral occipital* | 226 | July 17 2020 |
| **pFS** (posterior fusiform sulcus) | -36 | -45 | -18 | *objects* | 692 | May 14 2020 |
| **PPA** (parahippocampal place area) | -30 | -45 | -9 | *place* | 189 | Feb. 18 2021 |
| **SPOC** (superior parietal occipital cortex) | -18 | -78 | 39 | *reaching* | 99 | June 25 2019 |
| **aIPS** (anterior intraparietal area) | -42 | -33 | 45 | *grasping* | 90 | June 25 2019 |
| **PMv** (ventral premotor) | -56 | 7 | 31 | *grasping* | 90 | June 25 2019 |
| **PMd** (dorsal premotor) | -24 | -12 | 60 | *grasping* | 90 | June 25 2019 |
| **M1/S1** (primary sensory/motor) | -33 | -27 | 63 | *grasping* | 90 | June 25 2019 |

Primary visual cortex (V1) was included because it represents the first stage of cortical visual processing upon which all subsequent visuomotor computations rely. Primary motor area M1 was included instead as the final stage of processing, where motor commands are generated and sent to the arm and hand. In our study, however, we refer to this ROI as primary motor and somatosensory cortex M1/S1, because our volumetric data do not allow us to distinguish between the two banks of the central sulcus along which motor and somatosensory regions lie.

We next selected regions believed to perform the sensorimotor transformations that link the visual input to the motor output. The dorsal visual stream is thought to be predominantly specialized for visually guided actions, whereas the ventral stream mostly specializes in visual object recognition (Goodale and Milner, 1992; Culham et al., 2003; Cavina-Pratesi et al., 2007). Nevertheless, significant crosstalk occurs between these streams (Budisavljevic et al., 2018), and visual representations of object material properties have been found predominantly in ventral regions. We therefore selected areas across both dorsal and ventral visual streams that would encode grasp axis, grasp size, and object mass.

Regions that we expected could encode grasp axis were dorsal stream regions SPOC (Fattori et al., 2004, 2009, 2010; Monaco et al., 2011), aIPS (Taubert et al., 2010), PMv (Murata et al., 1997; Raos et al., 2006; Theys et al., 2012), and PMd (Raos et al., 2004). Regions that we expected could encode grasp size were dorsal stream regions SPOC, aIPS (Monaco et al., 2015), PMd (Monaco et al., 2015), and PMv (Murata et al., 1997; Raos et al., 2006; Theys et al., 2012), as well as ventral stream regions LO (Monaco et al., 2015). Finally, based on previous literature we expected visual estimates of object mass to be encoded in ventral stream regions LO, pFS, and PPA (Cant and Goodale, 2011; Hiramatsu et al., 2011; Gallivan et al., 2014; Goda et al., 2014, 2016). We further hypothesised that the network formed by aIPS, PMv, and PMd might play a role in linking ventral stream representations of object mass to the motor commands generated and sent to the hand through M1 (Murata et al., 2000; Borra et al., 2008; Janssen and Scherberger, 2015).

**Figure 1H** shows the locations of our selected ROIs as volumes within the Colin27 template brain. To locate all other left hemisphere ROIs (except V1) in a standardized fashion we searched the automated meta-analysis website *neurosynth*.*org* (Yarkoni et al., 2011) for key words (see Table 1), which resulted in volumetric statistical maps in *nifti* files. Visual inspection of the maps allowed us to locate the ROIs we had pre-selected based on a combination of activation peaks, anatomical criteria, and expected location from the relevant literature. For example, aIPS was selected based on the hotspot for “grasping” in Neurosynth nearest the intersection of the intraparietal and postcentral sulci (Culham et al., 2003). Spherical ROIs of 15-mm diameter, centred on the peak voxel, were selected for all regions except V1. Because Neurosynth is based on a meta-analysis of published studies, search terms like “V1” would be biased to the typical retinotopic locations employed in the literature and likely skewed towards the foveal representation (whereas the objects and hand would have been viewed across a larger expanse within the lower visual field). As such, we defined V1 in the left hemisphere’s V1 using the (Wang et al., 2015) atlas, which mapped retinotopic cortex +/-∼15° from the fovea. **Table 1** presents an overview of our ROI selection, where we list all our Neurosynth-extracted ROIs with their peak coordinates, search terms and download dates. We also share our ROIs (in MNI space) in the *nifti* format (doi upon acceptance).

#### Representational Similarity Analysis

The analysis of activation patterns within the selected ROIs was performed using multivoxel pattern analysis, specifically representational similarity analysis (RSA) (Kriegeskorte, 2008; Kriegeskorte et al., 2008). An activation pattern corresponded to the set of normalized β-weight estimates of the blood oxygenation level-dependent (BOLD) response of all voxels within a specific ROI for a specific condition. To construct representational dissimilarity matrices (RDMs) for each ROI, we computed the dissimilarity between activation patterns for each condition. Dissimilarity was defined as 1-r, where r was the Pearson correlation coefficient. RDMs were computed separately from both grasp planning and grasp execution phases. These neural RDMs computed were then correlated to model RDMs (**Figure 1D,E,F**) to test whether neural representations encoded grasp axis, grasp size, and object mass. To estimate maximum correlation values expected in each region given the between-participant variability, we computed the upper and lower bounds of the noise ceiling. The upper bound of the noise ceiling was computed as the average correlation of each participant’s RDMs with the average RDM in each ROI. The lower bound of the noise ceiling was computed by correlating each participant’s RDMs with the average of the other participants’ RDMs. All correlations were performed between upper triangular portions of the RDMs excluding the diagonal. We then used one-tailed Wilcoxon signed rank tests to determine whether these correlations were significantly >0 within each ROI. We set statistical significance at p<.05 and applied false discovery rate (FDR) correction for multiple comparisons following (Benjamini and Hochberg, 1995).

To visualize the representational structure of the neural activity patterns within grasp planning and grasp execution phases, we first averaged RDMs across participants in each ROI and phase. We then correlated average RDMs across ROIs within each phase, and used hierarchical clustering and multidimensional scaling to visualize representational similarities across brain regions. We also correlated average RDMs across ROIs and across planning and execution phases. Statistically significant correlations (p<.05 with Bonferroni correction) are shown also as topological connectivity plots (within-phase data) and as Sankey diagram (between-phase data).

#### Grasp Comfort Ratings

Grasp comfort ratings were analysed using simple t-tests to assess whether ratings varied across different grasp axes, grasp sizes, or object mass. The difference between ratings for each condition was then used to create grasp comfort RDMs for each participant. Grasp comfort RDMs were correlated to model RDMs to further test how strongly grasp comfort corresponded to grasp axis, grasp size, and object mass. To search for brain regions that might encode grasp comfort, the average grasp comfort RDM was correlated to neural RDMs following RSA as described above.

## Acknowledgments

We thank Mel Goodale for helpful discussions when designing the study.

